# Analysis of healthy reference ranges for clinically relevant gut microbial indicators: Mykinso cohort study in Japan

**DOI:** 10.1101/2020.02.25.965459

**Authors:** Satoshi Watanabe, Shoichiro Kameoka, Natsuko O. Shinozaki, Ryuichi Kubo, Akifumi Nishida, Minoru Kuriyama, Aya K. Takeda

## Abstract

**Background:** Although several large cohort studies have focused on the human gut microbiome, large cohort study of the Japanese gut microbiome is scarce, especially that of healthy or non-diseased individuals. The purpose of this study was to establish a reference range for gut microbial indices by collecting Japanese real-world microbiome data from a Mykinso cohort.

**Methods:** We collected stool samples and original survey on lifestyle from 5,843 Japanese people through Mykinso gut microbiome testing service. From the obtained 16S rRNA sequence data derived from the stool sample, the ratio and distribution of each taxon were analyzed. The relationship between different epidemiological attributes and gut microbial indicators were statistically tested.

**Results:** The qualitative and quantitative indicators of these common gut microbiota were confirmed to be strongly correlated with age, sex, constipation/diarrhea, and history of lifestyle-related diseases. Therefore, we set up the healthy sub-cohort that controlled for these attribute factors, and defined a reference range from the distribution of gut microbial index in that population.

**Conclusions:** The gut microbiota of Japanese people had high beta diversity, and there was no single “typical” gut microbiota type. We believe that the reference range of the gut microbial index obtained in this study can be a new reference value for determining the balance and health of the gut microbiota of an individual. In the future, it is necessary to clarify the clinical validity of this reference value by comparing with the clinical disease cohort.

## INTRODUCTION

Since the establishment of next-generation 16S rRNA sequencing analysis, multiple large cohort studies focused on the human gut microbiome have been conducted, such as the US Human Microbiome Project [1] and MetaHIT in Europe [2]. An integrated catalog of human fecal microbial metagenomes from 1200 people in the United States, China, and Europe has identified 9.9 million microbial genes across fecal microbiota [3]. However, the Japanese gut microbiome study is scarce, especially that of healthy or non-diseased individuals. Further, in recent studies comparing gut microbiomes by race/nationality, clear impacts of dietary habits were demonstrated [4], suggesting that, due to Japanese unique food culture, the gut of Japanese individuals could harbor different flora from individuals in western countries. Therefore, to advance research on the gut microbiome and various diseases in Japan, it is critical to characterize the healthy gut microbiome in the Japanese population.

Host parameters such as age, gender, and body mass index (BMI) have been reported to be related to individual differences in gut microbiota composition [5][6][7][8][9]. Further, differences in dietary habits have been shown to affect the bacterial diversity and enterotype of human gut microbiota [10][11], which may partially explain why differences in residential areas/countries are strongly associated with differences in gut microbiota composition [12][13].

Recently, several studies have revealed gender differences in gut microbiota [14][15][16][17]. Min et al [18] conducted an association study to identify bacterial compositions associated with men and women and showed similar microbiota characteristics, including overall abundance and diversity, between men and women. However, they also showed gender differences at the species level between microbial taxa related to fat distribution, suggesting the existence of a gender-specific microbiome signature corresponding to gender-specific fat distribution, which may also contribute to the observed sex-specific immunity differences [19]. Thus, several immune pathophysiology may be involved in gender differences in gut bacterial composition.

Age is also an important factor affecting the gut microbiota [8][20][21][22]. Recent reports have described differences in gut microbiota between children and adults, and an adult-like composition of bacterial communities is established around 3–4 years old or more [7] [8][23][24][25]. In addition, intestinal microbiota has been shown to change with aging, although the definition of old people differs by reports, such as over 60, 65, 70, or 100 [14][26][27][28]. However, the mechanism of change with aging is still unknown. Yatsunenko et al. [8] conducted a large study of subjects aged 0–83 years and showed continuous changes that occurred with age. Their report provided important insights. First, the period required to form an adult-like gut microbiota was the 3-year period following birth. Second, interpersonal variation was significantly greater between children than between adults. Third, the dominance of *Bifidobacterium* in the baby microbiota continued throughout the first year of life although this dominance composition diminished with age. Nevertheless, due to the limited number of subjects over 60 years of age, the continuous changes that occur in older people remain unknown. Recently, Odamaki et al. [7] reported age-related compositional differences from infants to centenarians in a Japanese cross-sectional study. They found that *Bifidobacterium* decreased and Enterobacteriaceae increased with age, as observed in some previous studies [8][20][21][22].

Relationships have also been shown between gut microbiota and diarrhea/constipation. Vandeputte et al. [29] described the association between stool consistency and gut microbiota composition in 53 healthy female subjects. Tigchelaar et al. [30] also reported an association between stool consistency and the structure of gut microbiota. Hadizadeh et al. [31] demonstrated a correlation between the number of bowel movements and gut microbiota. In a Japanese cohort, Takagi et al. [32] reported significant differences in microbial structure between individuals with different stool consistency (Bristol stool scale type). Therefore, investigating the relationship between bowel habits and intestinal bacterial composition can provide important information on gastrointestinal motility function.

However, Japan has its own food culture and customs compared to western countries, and the intestinal flora of Japanese individuals contain more genes for polysaccharide-degrading enzymes derived from water-soluble dietary fiber than Americans [33]. This feature may be related to the long life expectancy of Japanese people and the low body mass index (BMI) [34][35]. Nishijima et al. [36] clearly showed significant differences in the gut microbiota of the Japanese population compared to other countries, which cannot be explained by meals alone. Therefore, the structure of the intestinal flora may be highly dependent on an individual’s country/region and lifestyle [37]. In this study, we investigated the relative abundance ranges of these taxa in stool samples from a large healthy human cohort. Further, we analyze the relationship between the above-focused genera or the gut microbiota composition and the Japanese demographic features, lifestyle, and bowel habits. Finally, we developed a reference range using a large healthy Japanese cohort, and considered the effect of age, gender, diarrhea, and constipation.

## METHODS

### Study Design and Participants

From November 2015 through June 2019, a Mykinso cohort of 5,843 individuals who had submitted fecal samples (one sample per subject) were selected from the data obtained through the ongoing Mykinso gut microbiome testing service. Informed consent about the study was obtained from all participants. All procedures complied with the principles of the Declaration of Helsinki and were approved by the Institutional Review Board at our institution, registered as UMIN000028887 and UMIN000028888 in the UMIN Clinical Trials Registry System.

### Demographic features, bowel habits and disease and medication data

The metadata using original survey (Table S1) were collected through the Mykinso gut microbiome testing service. The original survey included questions on demographic features, lifestyle, bowel habits and disease. Individuals were scored positive for a disease if they replied yes to any original survey question, negative if they replied no, and unknown if data were unavailable across all original surveys.

### Fecal sampling, DNA extraction, and sequencing

Fecal samples were collected using brush-type collection kits containing guanidine thiocyanate solution (Techno Suruga Laboratory, Shizuoka, Japan), transported at normal temperature, and stored at 4 °C. DNA extraction from the fecal samples was performed using an automated DNA extraction machine (GENE PREP STAR PI-480, Kurabo Industries Ltd, Osaka, Japan) according to the manufacturer’s protocol. The V1–V2 region of the 16S rRNA gene was amplified using forward primer (16S_27Fmod: TCG TCG GCA GCG TCA GAT GTG TAT AAG AGA CAG AGR GTT TGA TYM TGG CTC AG) and reverse primer (16S_338R: GTC TCG TGG GCT CGG AGA TGT GTA TAA GAG ACA GTG CTG CCT CCC GTA GGA GT) with KAPA HiFi Hot Start Ready Mix (Roche). To sequence 16S amplicons by Illumina MiSeq platform, dual index adapters were attached using the Nextera XT Index kit. Each library was diluted to 5 ng/µL, and equal volumes of the libraries were mixed to 4 nM. The DNA concentration of the mixed libraries was quantified by qPCR with KAPA SYBR FAST qPCR Master mix (KK4601, KAPA Biosystems) using primer 1 (AAT GAT ACG GCG ACC ACC) and primer 2 (CAA GCA GAA GAC GGC ATA CGA). The library preparations were carried out according to 16S library preparation protocol of Illumina (Illumina, San Diego, CA, USA). Libraries were sequenced using the MiSeq Reagent Kit v2 (500 Cycles), 250 bp paired-end.

### Taxonomy assignment based on 16S rRNA gene sequences

The paired-end reads of the partial 16S rRNA gene sequences were clustered by 97% nucleotide identity, and then assigned taxonomic information using Greengenes database (v13.8)[38] through QIIME pipeline (v1.8.0)[39]. The steps for data processing and assignment based on the QIIME pipeline were as follows: (i) joining paired-end reads; (ii) quality filtering with an accuracy of Q30 (>99.9%) and a read length >300 bp; (iii)10,000 reads per sample were randomly extracted for the subsequent analysis; (iv) clustering operational taxonomic units (OTUs) with 97% identity by UCLUST (v1.2.22q) [40]; (v) assigning taxonomic information to each OTU using RDP classifier [41] with the full-length 16S gene data of Greengenes (v13.8) to determine the identity and composition of the bacterial genera.

### Transformation of compositional microbiome data for hypothesis testing

The centered log-ratio (clr) transformed values were used as inputs for multivariate hypothesis testing [42] to manage 0 count values as both point estimates using zCompositions R package [43] and as a probability distribution using ALDEx2 [44] available on Bioconductor.

### Group differences in beta-diversity

Aitchison distance, which is the Euclidean distance between samples after clr transformation, and the distances between samples are the same as the phylogenetic ilr [42]. The replacement for β-diversity exploration of microbiome data is the variance-based compositional principal component (PCA) biplot [45], in which the relationship between inter-OTU variance and sample distance can be observed [46]. Compositional PCA biplots display the relationships between OTUs and the distances between samples on a common plot to clean substantial qualitative information regarding the quality of the dataset and the relationships between groups [46].

### Group differences in alpha-diversity

Microbiota diversity was assessed by Shannon index based on 97% nucleotide sequence identity. These values were calculated by QIIME [[39]] with a depth of 10,000 reads. To test two-group differences between male and female groups, we calculated p-values by the two-sided unpaired Welch’s *t*-test. To test group differences among age-class group in the diversity index, we calculated *p* values by the one-way analysis of variance (ANOVA).

### Group differences in taxonomic abundance

To compare the taxonomic abundance between the groups, we conducted the univariate statistical test using the ALDEx2 tool [44]. The false discovery rate (FDR) control was performed based on the Benjamini-Hochberg procedure to correct for multiple testing, i.e., ‘p.adjust’ in R. Analysis was confined to taxa with a prevalence greater than 10% and a maximum proportion (relative abundance) greater than 0.005. An FDR-adjusted *p* value less than 5% was considered to be significant.

## RESULTS

### Cohort characteristics

Participants primarily resided in Japan (n = 5843) (Table S2), with a greater range in age, stool type, and lifestyle than other Japanese large-scale microbiome projects [7][32][47]. Using an original survey, participants (n = 4479) reported demographic features, disease history, and lifestyle data (participants missing any of these data were excluded, Table S3). In accordance with our institutional review board (IRB), all survey questions were optional (question response rate, 76.65%). Eligible subjects were male and female subjects who were considered to not have disease history (Ineligible subjects were those who self-reported any disease history (Table S4).). Eligible criteria included no self-reported history of any disease, including significant gastrointestinal inflammatory disease such as inflammatory bowel disease or functional gastrointestinal disorders such as irritable bowel syndrome (IBS). The primary investigation focused on a “non-disease adult” subset of individuals; no self-reported history of inflammatory bowel disease (IBD) and diabetes. Finally, 2865 individuals were included for subsequent analysis (Table 1, Supplementary Figure 1).

**Table 1.** Distribution of primary eligible subjects

| **Years** | **Female** | **Male** | **Number of**<br>**samples** |
| --- | --- | --- | --- |
| 19 under | 20 | 10 | 30 |
| 20-29 | 322 | 135 | 457 |
| 30-39 | 559 | 408 | 967 |
| 40-49 | 485 | 358 | 843 |
| 50-59 | 249 | 159 | 408 |
| 60-69 | 73 | 52 | 125 |
| 70 over | 14 | 21 | 35 |
| Sum | 1722 | 1143 | 2865 |

**Figure 1.**
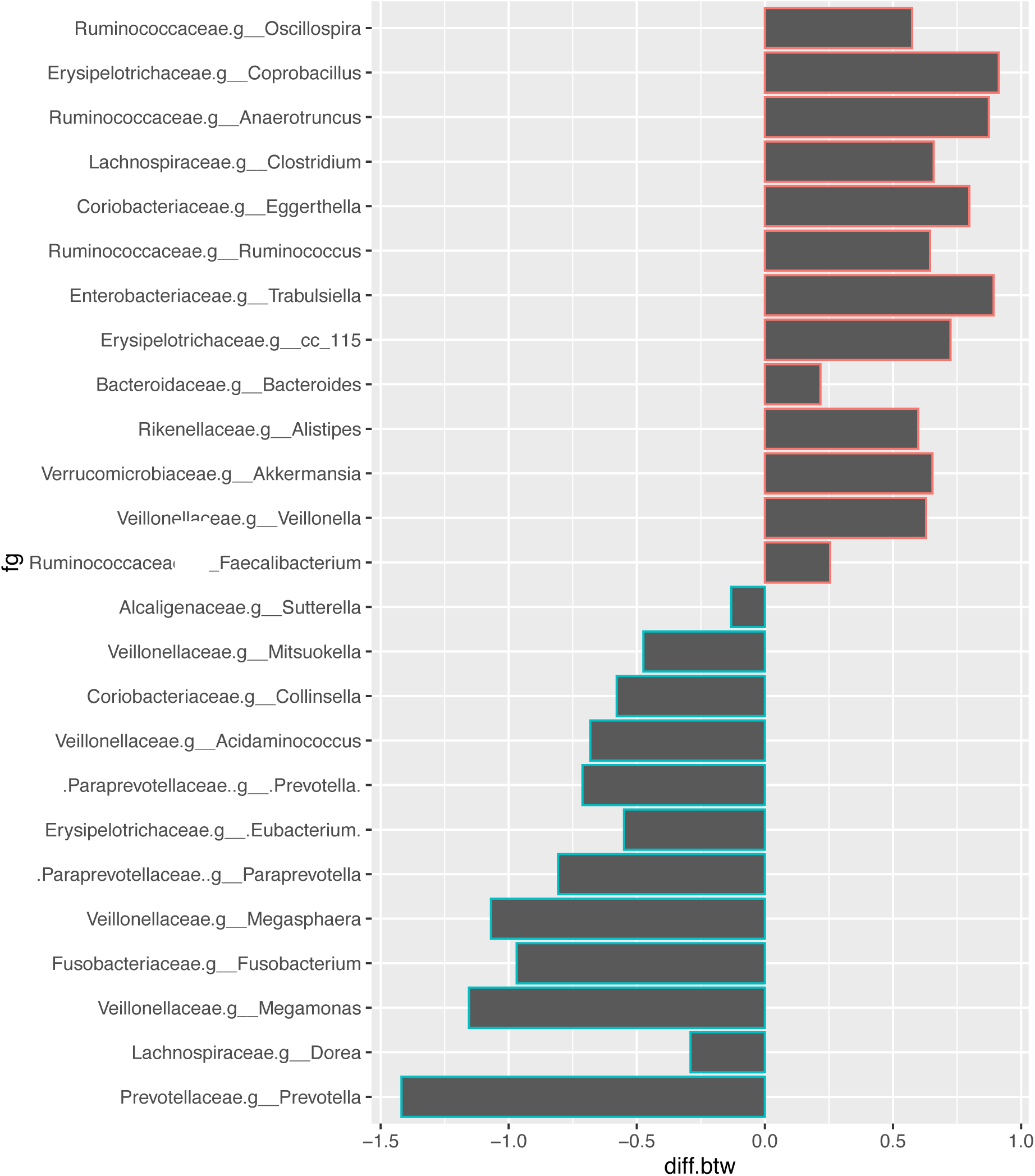
Relative abundance of gut microbiota in male and female subjects. Genera were significantly different between male and female subjects. diff_btw = median difference in clr values between female and male groups. magenta: positive diff_btw in female group; cyan: negative diff_btw in female group

### Sex-related gut microbiota

Taxonomic differences in the microbial community were evaluated at the genus level. As shown in Figure 1, the comparison of microbial composition between male and female subjects showed a significant richness in the abundance of 12 genera in male subjects and in 13 genera in female subjects (blue and red pch in Figure 1, respectively). These were characterized by a richness in representative genera *Prevotella, Megamonas, Collinsella, Dorea, Megasphaera*, and *Fusobacterium* (all *p* < 0.001) in male subjects and an increase in representative genera *Oscillospira, Coprobacillus, Ruminococcus, Bacteroides, Eggerthella, Anaerotruncus, Trabulsiella*, and *Akkermansia* (all *p* < 0.001) in female subjects. Subsequently, we evaluated the diversity of gut microbiota using the alpha-diversity index [Shannon index (OTU evenness estimation)]. The alpha-diversity index showed no statistically significant differences between male (mean = 6.013) and female (mean = 6.008) subjects (Welch Two Sample *t*-test; *t*(2352.5) = -0.198, *p* = 0.843, 95%CI = -0.049–0.060). Next, the overall structure of the gut microbiome for male and female subjects using beta-diversity indices was calculated for Aitchison distance (Figure 2). PCA revealed that there were microbial structural differences between male and female subjects (PERMANOVA, *R*^2^ = 0.060, *p* = 0.001).

**Figure 2.**
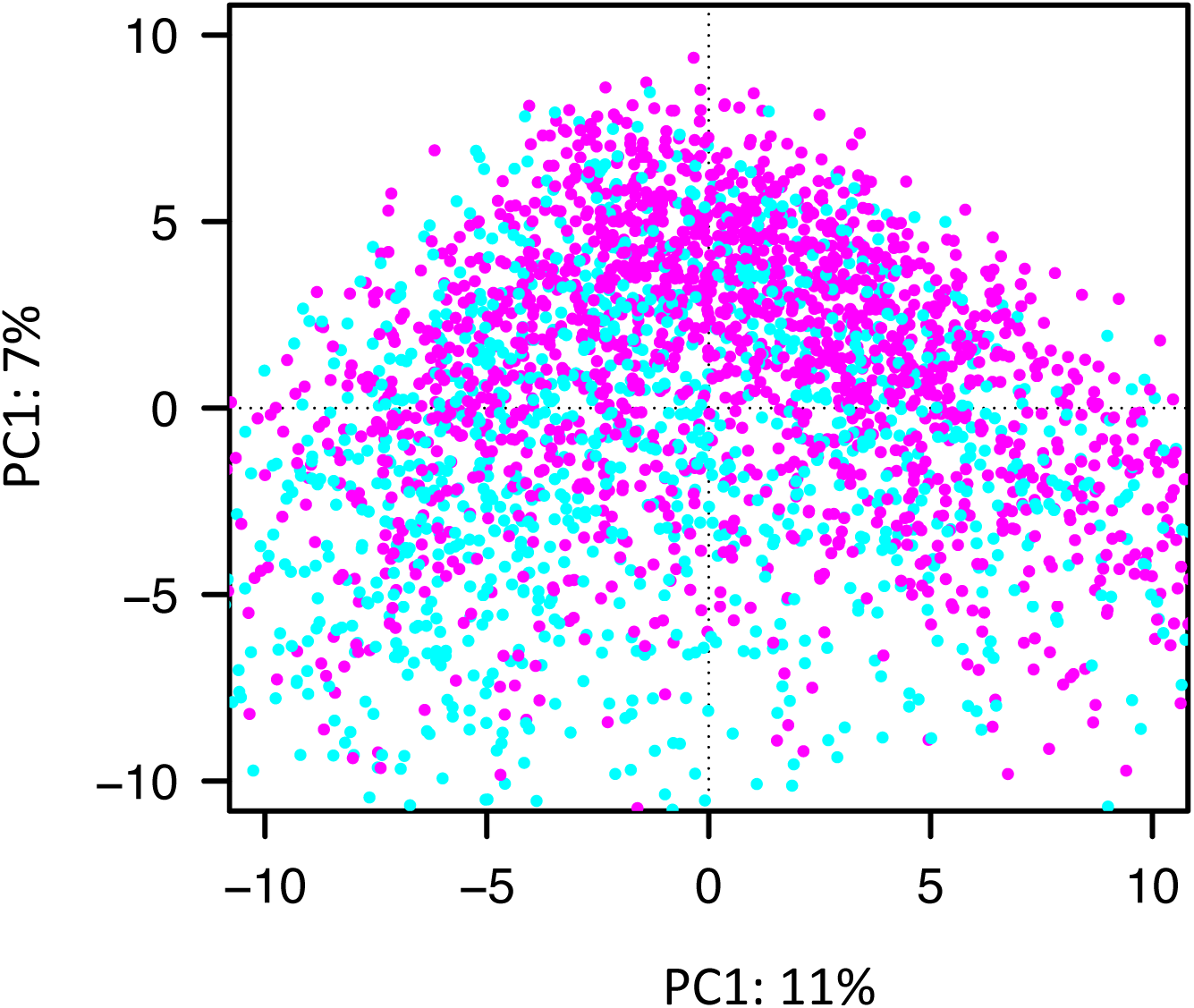
Plot of individual samples from PCA output (magenta: female samples, cyan: male samples). The distance between points is proportional to the Euclidian distance of CLR vectors of the samples (Aitchison distance). The multivariate distance between samples was estimated using the Aitchison distance, which showed significantly different composition between female and male samples (ANOSIM, *R*^*2*^ = 0.060, *p* = 0.001)

### Age-related gut microbiota

Further, differences in the gut microbial structure in each age group were taxonomically evaluated at the phylum level (Figure 3). In agreement with previous results, the microbiota composition included four predominant phyla (Firmicutes, Bacteroidetes, Actinobacteria, Proteobacteria). Of which, Actinobacteria showed significant decreases in 60 years subjects (*p* = 0.040, p.adjust = 0.640) and 70 years over subjects (*p* = 0.048, *p*.*adjus*t = 0.700) compared with 19 years under subjects.

**Figure 3.**
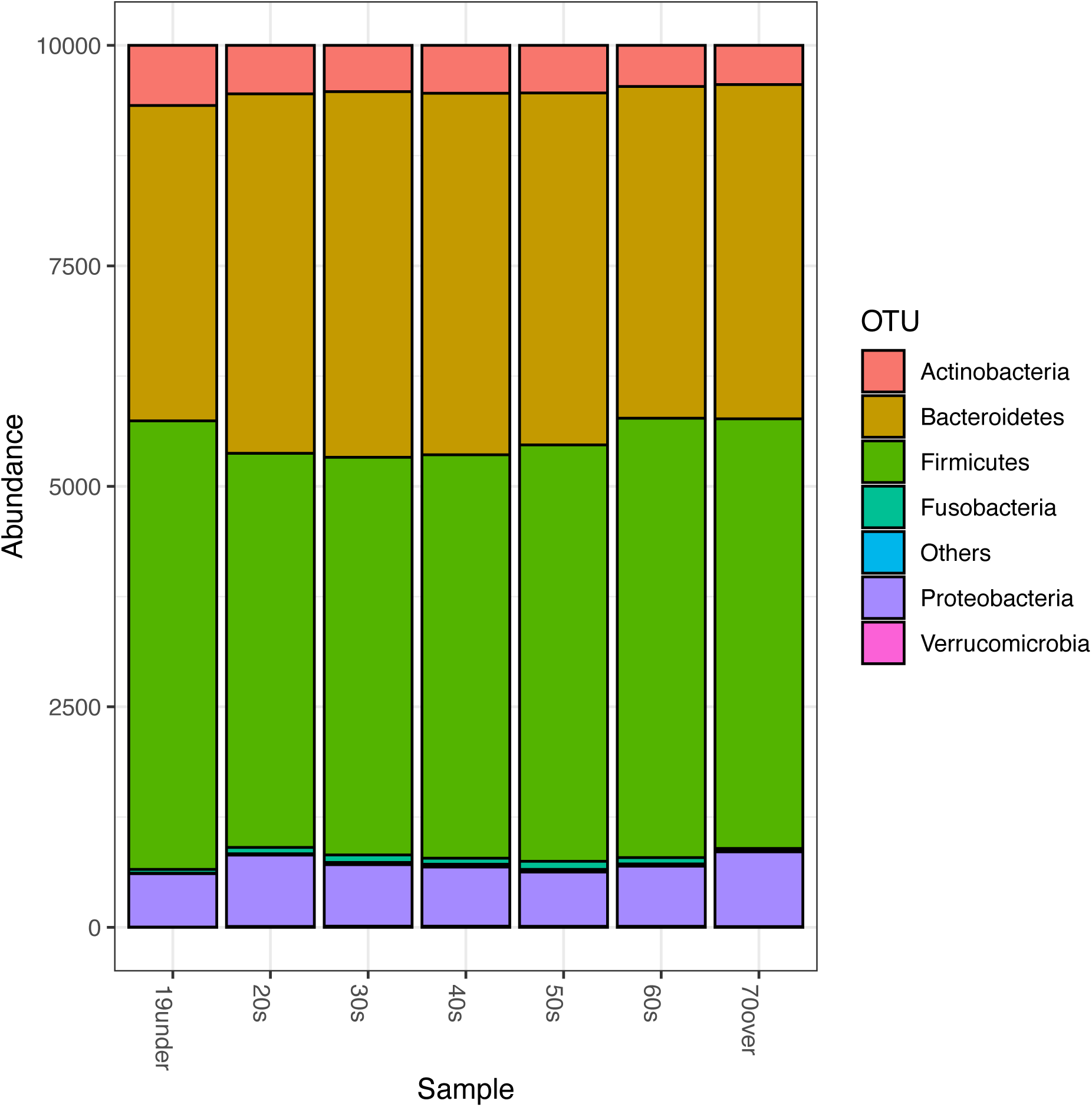
Comparative analyses of the taxonomic composition of the microbial community at the phylum level for each age group. Each component of the cumulative bar chart indicates a phylum

The alpha-diversity index showed significant differences across age groups in our cohort (ANOVA; *F*(6,2851) = 5.045, *p* < 0.001) (Figure 4). We also tested each pair difference with the Benjamini & Hochberg method, and statistically significant differences were found between 20s group and 60s group (*p* < 0.001), between 30s group and 60s group (*p* = 0.008), between 20s group and 40s group (*p* = 0.006), and between 50s group and 60s group (*p* = 0.043) (Table 2). Additionally, we created three age groups (0-19, 20–59, and 60 years or older), and the overall structure of the gut microbiome using beta-diversity indices was calculated for Aitchison distance (Figure 5). PCA revealed that there were microbial structural differences among the three age groups (PERMANOVA, *R*^2^ = 0.034, *p* < 0.001).

**Table 2.** Pairwise comparisons of Shannon-index between age groups

| **pair** | **diff** | **lwr** | **upr** | **p.adj** |
| --- | --- | --- | --- | --- |
| 20-19under | 0.001 | -0.398 | 0.400 | 1.000 |
| 30-19under | 0.109 | -0.284 | 0.502 | 0.983 |
| 30-20 | 0.108 | -0.012 | 0.228 | 0.112 |
| 40-19under | 0.151 | -0.242 | 0.545 | 0.917 |
| 40-20 | 0.151 | 0.027 | 0.274 | 0.006 |
| 40-30 | 0.043 | -0.057 | 0.142 | 0.870 |
| 50-19under | 0.128 | -0.272 | 0.529 | 0.965 |
| 50-20 | 0.127 | -0.017 | 0.272 | 0.125 |
| 50-30 | 0.019 | -0.106 | 0.145 | 0.999 |
| 50-40 | -0.023 | -0.151 | 0.105 | 0.998 |
| 60-19under | 0.349 | -0.082 | 0.779 | 0.203 |
| 60-20 | 0.348 | 0.134 | 0.562 | <0.001 |
| 60-30 | 0.240 | 0.039 | 0.441 | 0.008 |
| 60-40 | 0.197 | -0.006 | 0.400 | 0.063 |
| 60-50 | 0.220 | 0.004 | 0.437 | 0.043 |
| 70over-19under | 0.303 | -0.224 | 0.830 | 0.618 |
| 70over-20 | 0.302 | -0.069 | 0.674 | 0.199 |
| 70over-30 | 0.194 | -0.170 | 0.559 | 0.701 |
| 70over-40 | 0.152 | -0.214 | 0.517 | 0.885 |
| 70over-50 | 0.175 | -0.198 | 0.548 | 0.812 |
| 70over-60 | -0.046 | -0.451 | 0.359 | 1.000 |

diff: Differences in mean levels
lwr: 95% confidence lower level
upr: 95% confidence upper level
p.adj: p-value adjusted by Benjamin & Hochberg methods

**Figure 4.**
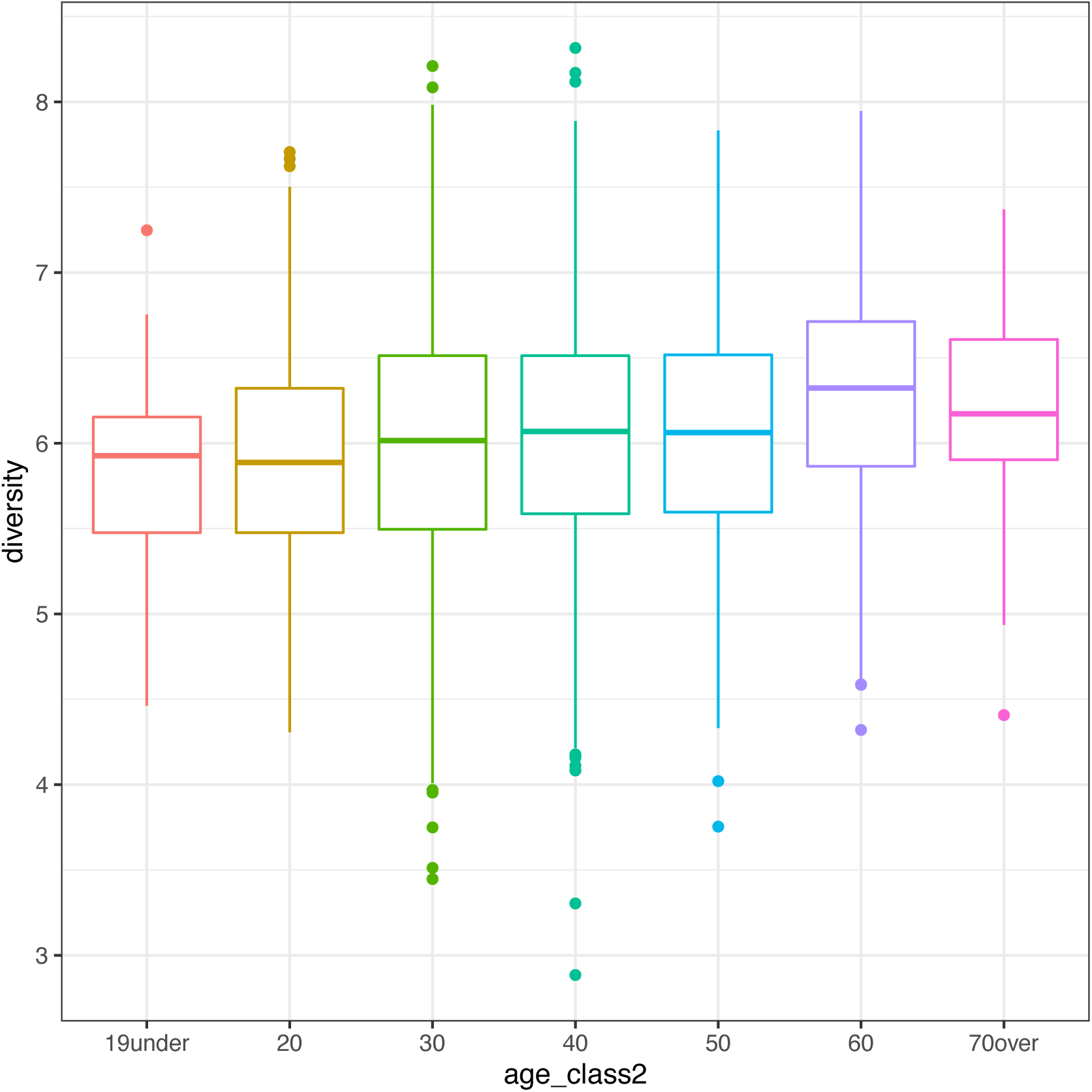
Age-related difference in alpha-diversities of gut microbiota

**Figure 5.**
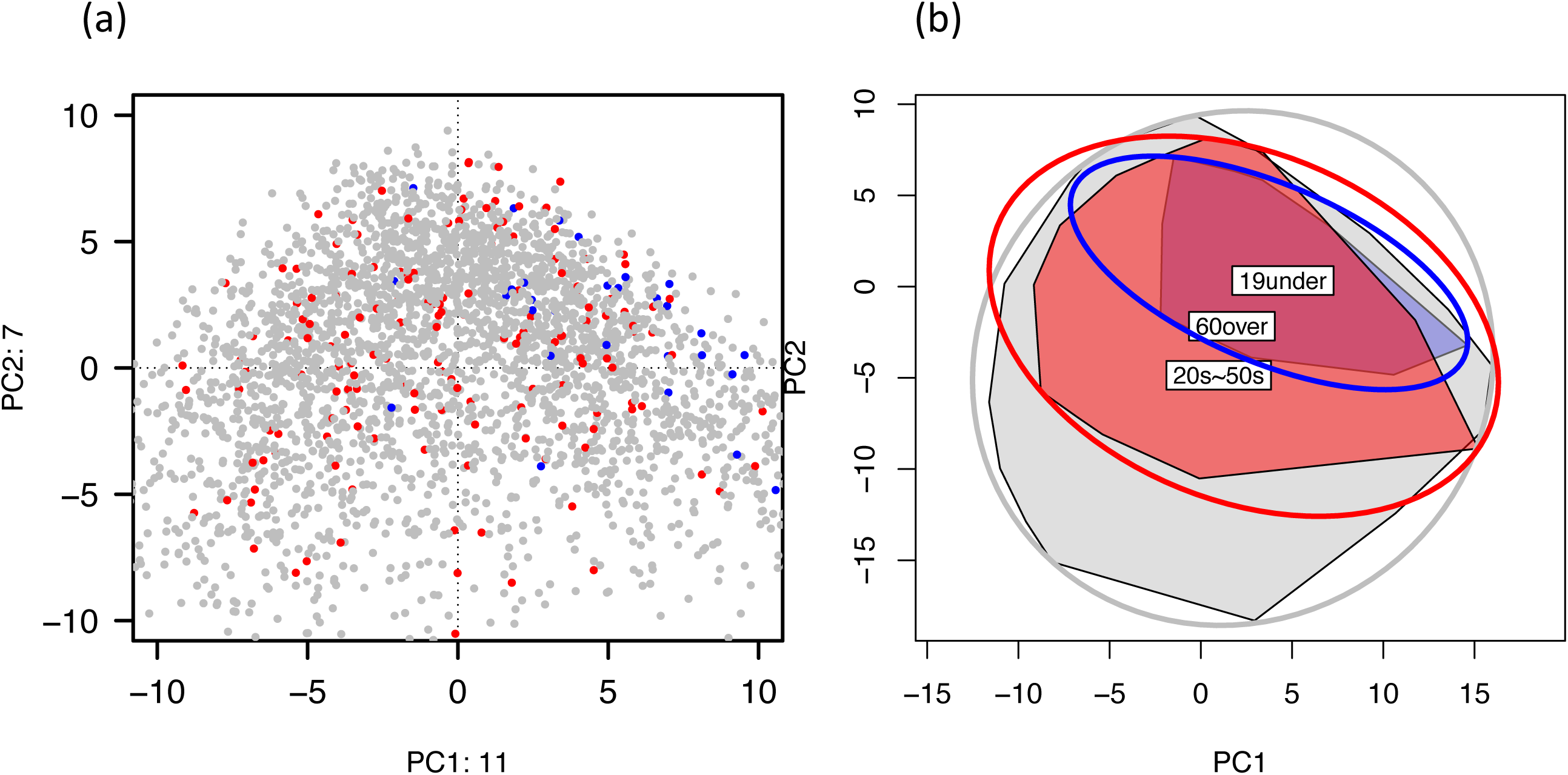
(a) Plot of individual samples from PCA output (red: senior samples, blue: junior samples, and gray: adult samples). The distance between points is proportional to the Euclidian distance of CLR vectors of the samples (Aitchison distance). (b) The multivariate distance between samples was estimated using the Aitchison distance, which showed significantly different composition in the junior, adult and senior samples (red: senior, blue: junior, and gray: adult) (PERMANOVA, *R*^2^ = 0.034, *p* < 0.001)

### Bowel habits-related gut microbiota

Considering the heterogeneity and varying generations of samples in this dataset, we excluded samples from children (0–19 years) and aged adults (over 60), which might cause bias in the subsequent analysis [14][26][27][28][7]. As a result, 2675 samples were included in the resulting dataset (Table 3, Supplementary Figure 1).

**Table 3.**
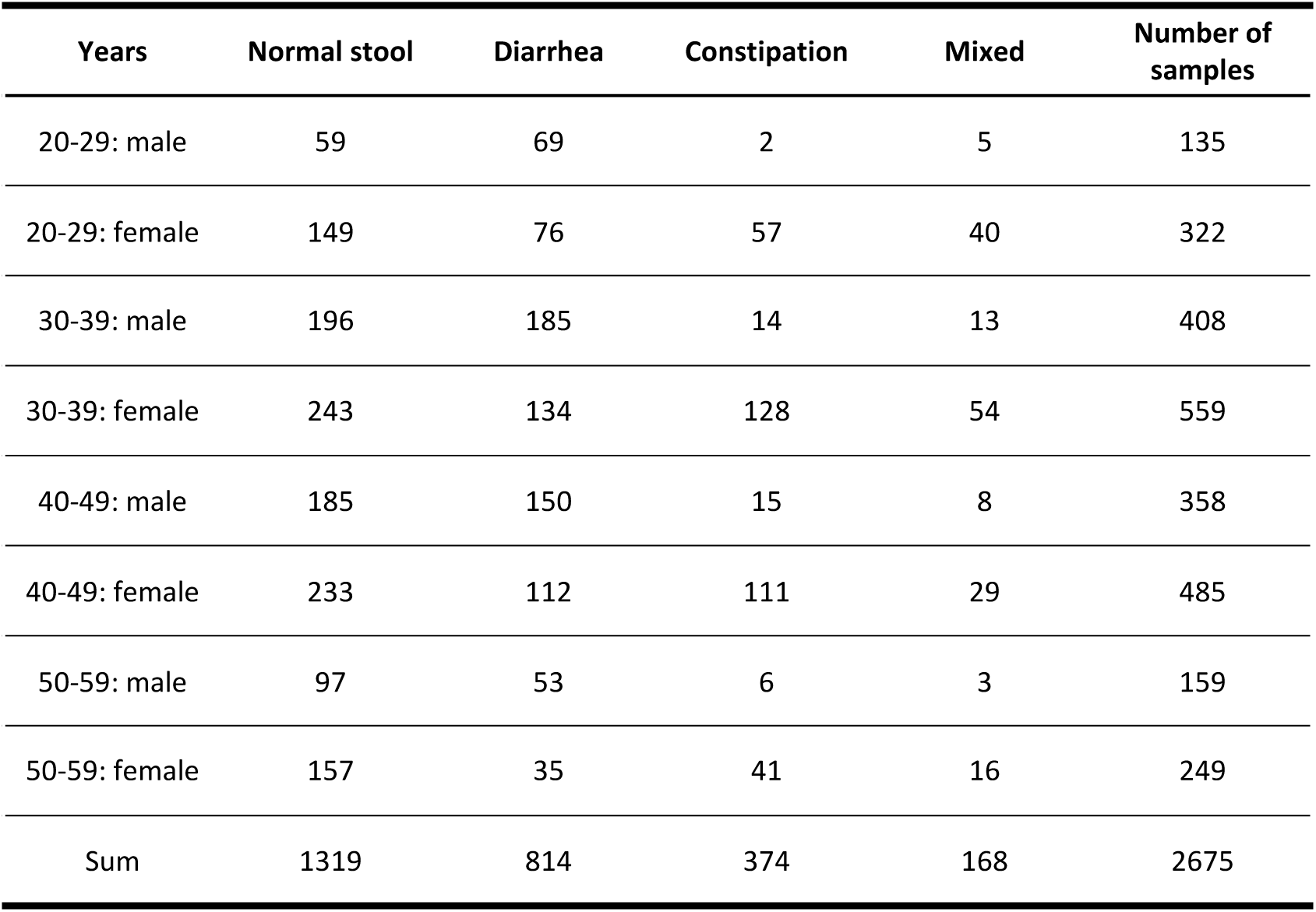
Distribution of bowel habits by sex and age-class (20–50)

| Years | Normal stool | Diarrhea | Constipation | Mixed | Number of samples |
| --- | --- | --- | --- | --- | --- |
| 20-29: male | 59 | 69 | 2 | 5 | 135 |
| 20-29: female | 149 | 76 | 57 | 40 | 322 |
| 30-39: male | 196 | 185 | 14 | 13 | 408 |
| 30-39: female | 243 | 134 | 128 | 54 | 559 |
| 40-49: male | 185 | 150 | 15 | 8 | 358 |
| 40-49: female | 233 | 112 | 111 | 29 | 485 |
| 50-59: male | 97 | 53 | 6 | 3 | 159 |
| 50-59: female | 157 | 35 | 41 | 16 | 249 |
| Sum | 1319 | 814 | 374 | 168 | 2675 |

The bowel habits (stool shape and defecation frequency) of all participants enrolled in this study were recorded and classified using the self-reported original survey. Additionally, perceived symptoms of diarrhea/constipation were recorded and classified. According to the stool shape, bowel frequency, and perceived symptom scores, participants were classified as normal bowel habit type, diarrhea type, constipation type, or mixed type. For the constipation type, when stool types 1 (hard stool), defecation frequency type 4 (less one time per week), or perceived constipation symptoms frequented within 1 month, they were classified as constipation. For the diarrhea type, when stool types 7 (liquid stools), defecation frequency type 1 (more three times per day), or perceived diarrhea symptoms frequented within 1 month, they were classified as diarrhea. And when the patient had both the constipation type and the diarrhea type, they were classified as mixed type. Constipation group (female, n = 337, 20.87%; male, n=37, 3.49%), diarrhea group (female, n =357, 22.11%; male, n = 457, 43.11%), mixed group (female, n =139, 8.61%; male, n = 29, 2.74%), and normal group (female, n = 782, 48.42%; male, n = 537, 50.66%) were observed (Table 3). Importantly, the alpha-diversity index for each bowel habit group showed a significant difference among groups in our cohort (ANOVA; *F*(3,2667) = 1.761, *p* < 0.001) (Figure 6). We also tested each pair difference with the Benjamini & Hochberg method, which showed statistically significant differences between the normal and diarrhea groups (*p* < 0.001), constipation and diarrhea groups (*p* < 0.001), and mixed and diarrhea groups (*p* = 0.001). These differences are visualized in Figure 6. Additionally, the overall structure of the gut microbiome among the three bowel habit groups using beta-diversity indices was calculated for Aitchison distance and visualized by PCA according to Aitchison distance (Figure 7). An additional PERMANOVA analysis based on categorical variables of their abundance showed that the bowel habit type was a significant factor contributing to the variation of the gut microbiota (*p* < 0.001). Approximately 0.7% of the variance in beta diversity was explained by the bowel habit type (PERMANOVA; *F*(3,2667) = 6.714, *R*^2^ = 0.007, *p* < 0.001), which was competitive with many microbiome covariates.

**Figure 6.**
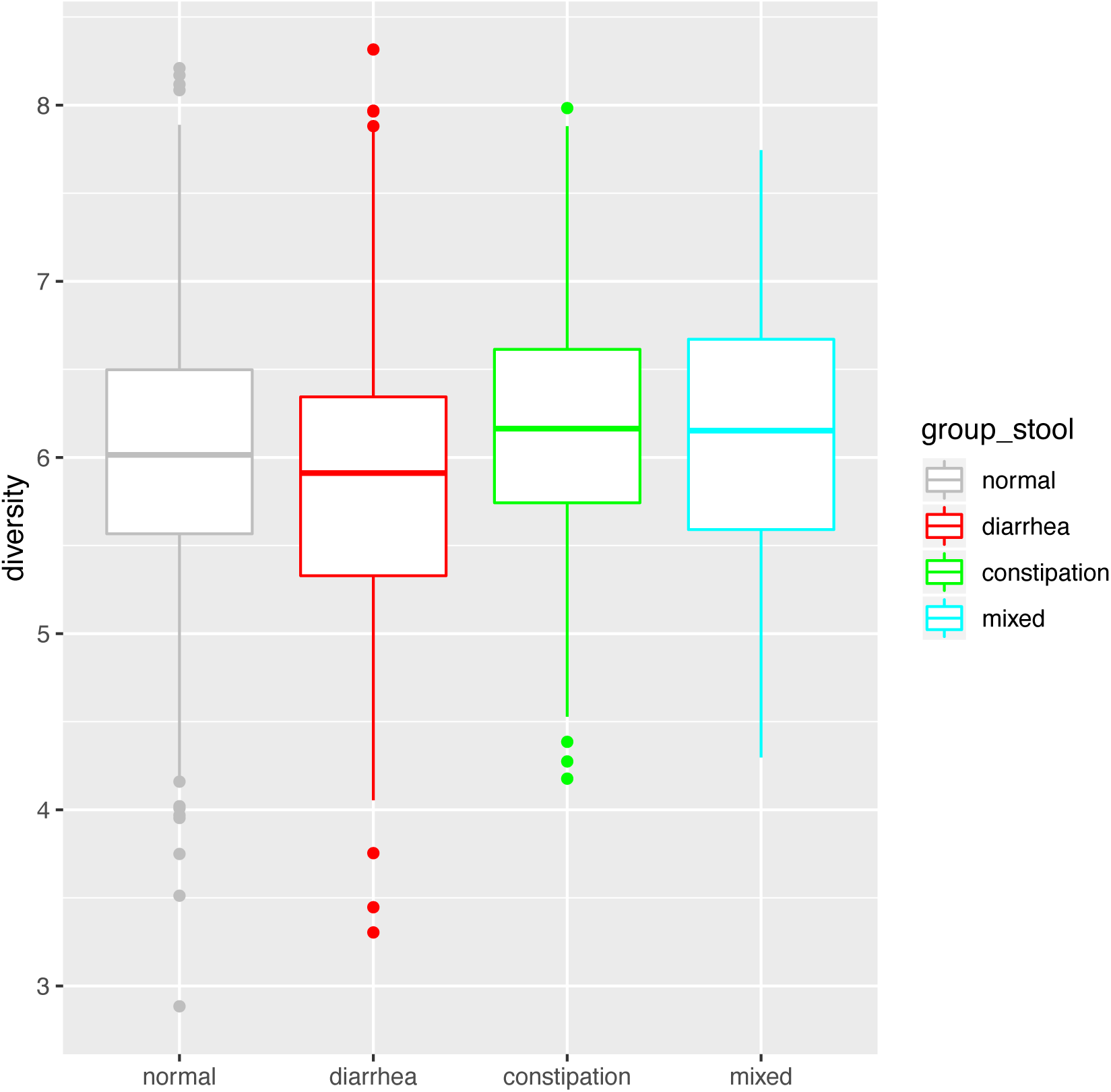
Alteration of alpha-diversity of gut microbiota associated with stool consistency. Comparison of α-diversity indices: Shannon-index (OTU evenness estimation). Bowel habit was categorized into three groups: normal, diarrhea, and constipation

**Figure 7.**
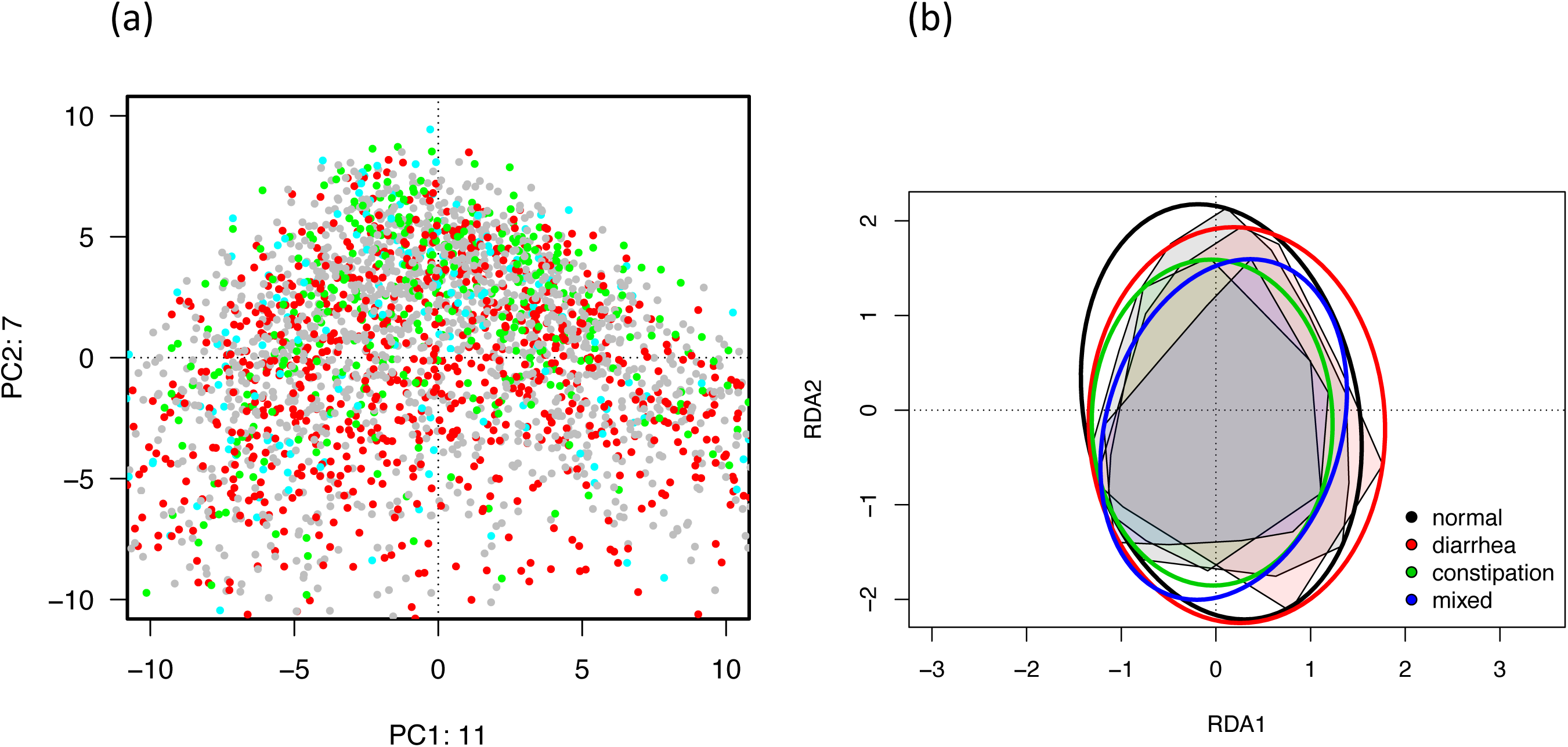
(a) Plot of individual samples from PCA output (gray: normal samples, red: diarrhea samples, green: constipation samples, and blue: mixed-type samples). The distance between points is proportional to the Euclidian distance of CLR vectors of the samples (Aitchison distance). The multivariate distance between samples was estimated using the Aitchison distance, which showed significantly different compositions in the junior, adult and senior samples (PERMANOVA, *R*^2^ = 0.007, *p* < 0.001). (b) RDA triplot of CLR vectors of the samples constrained by bowel habit group

Subsequently, at the genus level, we identified several altered bacteria among the three bowel habit groups. Interestingly, a significantly higher relative abundance of *Fusobacterium* in the diarrhea group and *Oscillospira* (*p* < 0.001) in the constipation group was observed. In addition, the relative abundance of *Ruminococcus, Anaerotruncus, Alistipes*, and *Akkermansia* (all *ps* < 0.001) were significantly higher in the constipation group, while *Dorea* (p < 0.01) was higher in the diarrhea group.

### Reference ranges from Japanese healthy cohort

Considering the heterogeneity and bowel habits of the sample dataset, we excluded samples from individuals with diarrhea or constipation (Supplementary Figure 1), which might cause bias in the reference range [29][30][31][32]. Finally, 1319 samples were selected as the healthy reference dataset (Supplementary Figure 1). We identified 453 genera and 20 phyla of Bacteria and Archaea in the gut microbiomes of the healthy reference dataset. The genera with an average relative abundance of ≥ 0.5% in the Japanese healthy reference dataset are listed in Supplementary Figure 2. At the genus level, the Japanese healthy reference was characterized by the highest abundance of *Bacteroides, Faecalibacterium, Prevotella, Blautia, Bifidobacterium, Coprococcus*, and *Parabacteroides* (Supplementary Fig 2).

In this study, health-related microbiome indices were selected based on peer-reviewed studies in academic journals and in-house data analyses (Table 4). To determine the reference range of 11 target microbiome indices, the dataset of 1319 individuals selected by the above definition from the Mykinso cohort was established. Microbiome data from this dataset were analyzed to determine the empirical reference ranges for two indices of overall community structure, two complex genus indices, one class, and six genera. For each of the 1319 samples, we determined the relative abundance of each target within the microbial population, revealing the distribution of the relative abundance of each target in the cohort (Table 4). These data were used to define a central 80% healthy range with confidence intervals for each target. Many of the targets show the significant spread, highlighting the importance of defining a reference range for health-related genera.

**Table 4.** Reference ranges from healthy Japanese subjects for 11 clinically relevant indices

| Microbiome Index | unit | group | lower limit<br>[10%] | Median<br>[50%] | upper limit<br>[90%] | Reference |
| --- | --- | --- | --- | --- | --- | --- |
| Shannon | value | all | 5.08 | 6.01 | 6.88 | [53, 54] |
|  |  | male | 5.15 | 6.03 | 6.93 |  |
|  |  | female | 5.05 | 6 | 6.85 |  |
| Observed genera | genus | all | 54 | 65 | 80 | [53, 54] |
|  |  | male | 54 | 65 | 81 |  |
|  |  | female | 54 | 66 | 79 |  |
| Bifidobacterium | % | all | 0.18 | 2.47 | 9.6 | [55, 56] |
|  |  | male | 0.12 | 2.18 | 8.45 |  |
|  |  | female | 0.21 | 2.79 | 10.41 |  |
| Faecalibacterium | % | all | 0.55 | 6.83 | 12.87 | [57] |
|  |  | male | 0.37 | 6.36 | 12.09 |  |
|  |  | female | 0.57 | 7.23 | 13.27 |  |
| butyrate-producing<br>genus | % | all | 4.25 | 12.16 | 20.47 | [54] |
|  |  | male | 3.64 | 11.53 | 20.03 |  |
|  |  | female | 4.66 | 12.88 | 20.89 |  |
| Clostridium | % | all | 0 | 0.19 | 0.79 | [54] |
|  |  | male | 0 | 0.19 | 0.7 |  |
|  |  | female | 0 | 0.2 | 0.89 |  |
| lactic acid-producing<br>genus | % | all | 0 | 0.01 | 0.22 | [54, 58] |
|  |  | male | 0 | 0.01 | 0.24 |  |
|  |  | female | 0 | 0.01 | 0.21 |  |
| Streptococcus | % | all | 0.05 | 0.38 | 2.58 | [54, 58] |
|  |  | male | 0.03 | 0.34 | 2.46 |  |
|  |  | female | 0.05 | 0.41 | 2.65 |  |
| Oral bacteria related<br>genus | % | all | 0.18 | 1.4 | 9.74 | [59, 60] |
|  |  | male | 0.15 | 1.31 | 9.71 |  |
|  |  | female | 0.2 | 1.48 | 9.9 |  |
| Fusobacterium | % | all | 0 | 0 | 1.37 | [59, 60] |
|  |  | male | 0 | 0 | 2.92 |  |
|  |  | female | 0 | 0 | 0.83 |  |
| Observed genera within<br>class gamma<br>proteobacteria | class | all | 2 | 5 | 8 | [60] |
|  |  | male | 2 | 5 | 8 |  |
|  |  | female | 2 | 5 | 8 |  |

## DISCUSSION

We developed a reference range using a large healthy Japanese cohort. The reference range considers the effect of age, gender, diarrhea, and constipation to aid physicians with accurate diagnosis of the intestinal bacterial composition ratio using a standard value derived from a healthy population. Eighteen intestinal bacterial indicators suggested to be associated with health status were selected. Using intestinal bacterial composition test panels, the detection of intestinal bacterial indicators outside the healthy range can be useful to reinforce the medical plan.

### Gut microbiota and sex/gender

Some characteristics of gender-specific immune differences are induced by gut microbiota. Fransen et al [48] investigated significant differences in bacterial groups at family or genus levels. Females had a higher abundance of *Desulfovibrionaceae, Lactobacillaceae* (*Lactobacillus* at the genus level), and *Verrucomicrobiaceae* (*Akkermansia* at the genus level), whereas males had a higher abundance of *Ruminococcaceae* and *Rikenellaceae* (*Alistipes* at the genus level). In this study, several characteristic differences were observed between male and female subjects regarding the genus abundance of gut microbiota. The genera *Prevotella, Megamonas, Fusobacterium*, and *Megasphaera* were significantly rich in male subjects, while *Bifidobacterium, Ruminococcus*, and *Akkermansia* were significantly rich in female subjects. These results are consistent with the results of previous Japanese studies [7][32], and may be considered as the characteristic of gender differences in the composition of intestinal microbiota in the Japanese population.

### Gut microbiota and age/generation

Recent reports have shown a clear difference in the composition of the intestinal microbiota of infants, adults, and the elderly [7][8][32]. The microbiota composition initially shifts after birth, followed by significant shifts during childhood and in later years [7][8]. In this study, we segmented by age group (young group, middle-aged group, elderly group). The diversity index showed slight differences between each age group from 20 to 89 years old (Figure 4), which are consistent with previous reports [7]. Further, our results are in agreement with studies indicating clear differences in gut microbiota composition among infants, adults, and the elderly [7][49][50]. Moreover, this study indicated that Actinobacteria abundance and alpha-diversity index were gut microbiome indices related to aging.

### Gut microbiota and bowel habits (diarrhea/constipation)

In this study, we evaluated the association between bowel habits and gut microbiota. Similar to Vandeputte et al. [29], we found a significant association between bowel habits (stool shape and defecation frequency) and gut microbiota diversity. Furthermore, deviation of the gut microbiota composition in several genera, including *Oscillospira, Ruminococcus, Anaerotruncus, Alistipes*, and *Akkermansia*, in constipation subjects and *Fusobacterium* and *Dorea* in diarrhea subjects were confirmed, which are consistent with a previous report [32]. Although the role of these genera in stool consistency remains unclear, the results illustrate the effect of gut microbiota on stool consistency in Japanese healthy subjects.

### Gut microbiota and racial/regional differences

Our results showed that Japanese adults (20–59 years old) had a greater abundance of genera *Bacteroides* and *Faecalibacterium* (interquartile ranges (IQR) of 27.43% (19.03–35.26) and 6.83% (3.39–10.06), respectively) and a relatively lower abundance of genera *Clostridium* (IQR 0.20 (0.04–0.44) %), compared to previous studies in other Japanese cohorts [36]. However, the estimated abundance of *Bifidobacterium* and *Blautia* was greater (interquartile ranges (IQR) of 2.47% (0.86–5.78) and 5.31% (2.94-7.85), than a previous study in other nations, which were < 0.5% and > 5%, in the US and China, respectively [36]. These bacterial compositions may be characteristic of the intestinal microbiota of the Japanese population, but may also reflect differences in DNA extraction methods [51][52] and the amplified region of the 16S rRNA [53]. The high abundance of *Bifidobacterium* has been also observed in the gut microbiome of Japanese children based on the 16S rRNA gene analysis [12], indicating its high prevalence throughout the Japanese population. *Bifidobacterium* is thought to be a beneficial microbe that contains more glycoside hydrolases for degrading starch than other intestinal microbes [54][55]. Therefore, the high abundance of *Bifidobacterium* may be the consequence of the intake of various saccharides in traditional and unique Japanese foods. However, it is unknown which foods or nutrients contribute to the high abundance of *Bifidobacterium*. As future prospects, it is essential to create a reference microbiota for the Japanese population by age group, increasing the number of subjects in the young and the elderly age groups. Additionally, investigations of geographical differences within Japan are of interest.

### Clinical relevance and Reference ranges

All 11 microbiome indices successfully identified using 16S rRNA gene sequencing were associated with specific health conditions. The alpha-diversity, including the Shannon index, and observed genera number appeared to be associated with better health [56]. A recent meta-analysis has proposed reduced alpha diversity as a reliable indicator of diarrhea-associated dysbiosis [57]. These microbiota diversity indices (Shannon index and observed OTUs) showed significant differences in the healthy aging group, indicating that a healthy, diverse diet promotes greater diversity in gut microbiota [56].

Previous studies have proposed that *Bifidobacterium* is inversely associated with IBD and diarrhea-associated dysbiosis [58], and the consumption of probiotics, inulin, and oligofructoses promotes an increase in *Bifidobacterium* abundance [59]. Additionally, *Faecalibacterium* has been proposed as a dominant member of human intestinal microbiota in healthy adults, and especially as a health sensor for active Crohn’s disease patients [60]. A recent meta-analysis showed a reduction in butyrate-producing *Clostridiales*, including *Coprococcus, Roseburia, Butyricicoccus, Faecalibacterium, Anaerostipes*, and *Butyricimonas*, which have been associated with a healthy gut [57].

While not all genera within the order *Lactobacillales* are verified lactic acid producers, the dominant genera within this order (including *Lactobacillus, Pediococcus, Leuconostoc, Lactococcus, Weissella*) are known to harbor genes for lactic acid production and are often enriched in case patients across multiple diseases [57]. *Lactobacillales* genera have been shown to adapt to the lower pH of the upper gastrointestinal tract [61]. Thus, the shared disease-associated taxa may be indicators of shorter stool transit times and disruptions in the redox state and/or pH of the lower intestine, rather than specific pathogens. Genera within *Lactobacillaceae* and *Streptococcaceae* families are dominant in the upper gastrointestinal tract and are present in the stool of many individuals at low frequency [57]. These taxa likely become enriched with faster stool transit time (i.e., diarrhea signatures) [57].

Previous studies have proposed Fusobacterium to be associated with various human diseases [62]. Dysbiosis associated with colorectal cancer is generally characterized by increased prevalence of known pathogenic or pathogen-associated *Fusobacterium* and *Enterobacter* genera, which were higher in colorectal cancer patients in two or more studies [57]. Furthermore, other oral community genera, such as members of *Porphyromonas, Peptostreptococcus*, and *Parvimonas*, were found with *Fusobacterium* on colonic tumors [63].

Regardless of health status, there are many microorganisms that are clinically related to health and disease in the intestines of all people, and the exquisite balance of these microorganisms varies greatly from person to person, making the definition of “good flora” difficult. However, in order to understand and monitor the health and balance of an individual’s gut microbiota, it is essential to first know the reference range available from a large, healthy population such as the one presented here. By further expanding the cohort of healthy subjects and accumulating a cohort of various health conditions and attributes, more valuable indicators can be identified, leading to the realization of personalized precision medicine using microbiome information in the future.

## Acknowledgments

We thank Dr. Shota Nakamura (Osaka University), iMet Laboratory team (Osaka University) and Biken Biomics, Inc. for sequencing and technical support, and Dr. Takeaki Uno (National Institute of Informatics) for support with data analysis. We also would like to thank Editage (www.editage.com) for English language editing.

## TABLES

**Table S1** Original survey questions

**Table S2** Distribution of all subjects

**Table S3** Distribution of question non-responding subjects

**Table S4** Distribution of subjects excluded from reference population due to reporting at least one of disease history

