## Supplemental Figures for "Analysis of healthy reference ranges for clinically relevant gut microbial indicators: Mykinso cohort study in Japan"

Title: Analysis of healthy reference ranges for clinically relevant gut microbial taxa in a health Japanese cohort
Journal: Journal of Gastroenterology
Authors: Satoshi Watanabe, Shoichiro Kameoka, Natsuko Shinozaki, Ryuichi Kubo, Akifumi Nishida, Minoru Kuriyama, , Aya K. Takeda
Affiliation: Cykinso, Inc, Tokyo, Japan

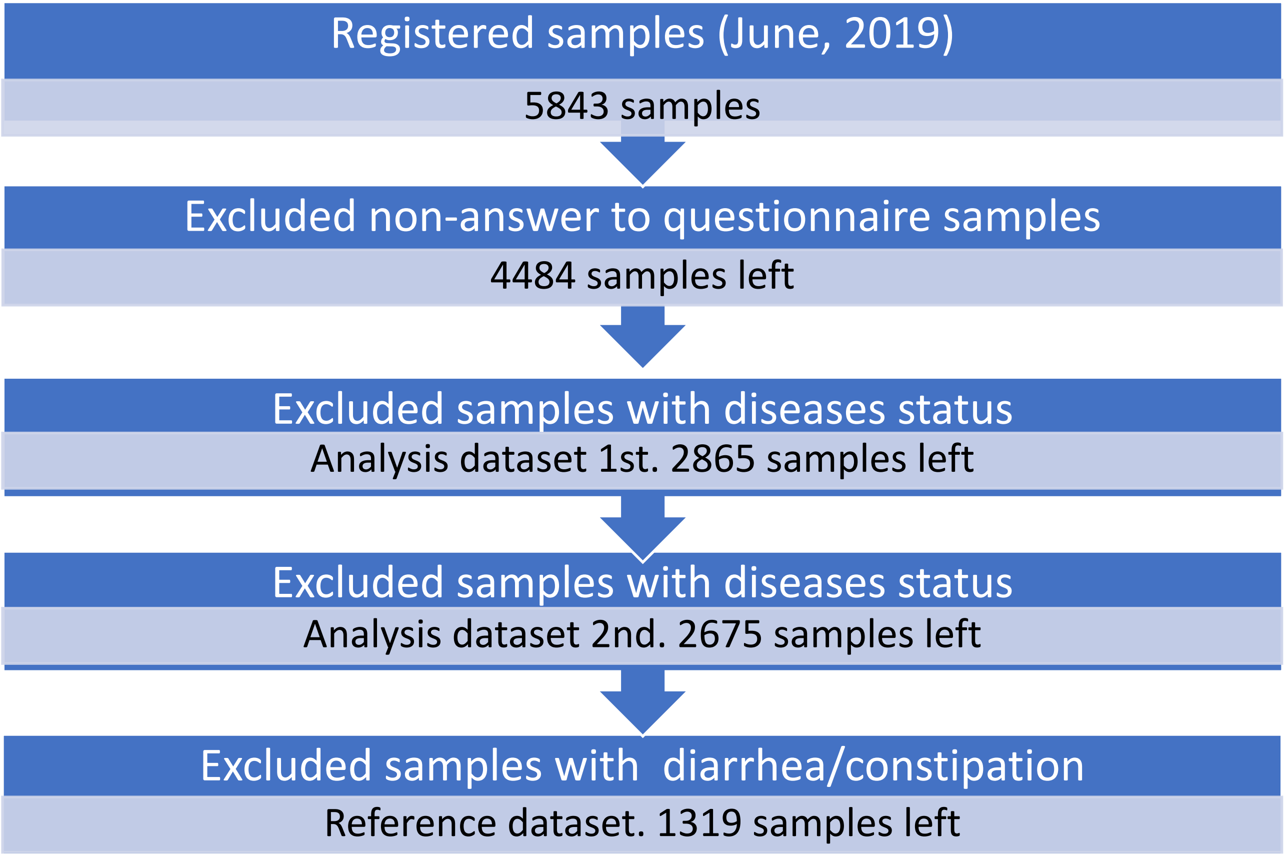


- Supplementary Figure 1

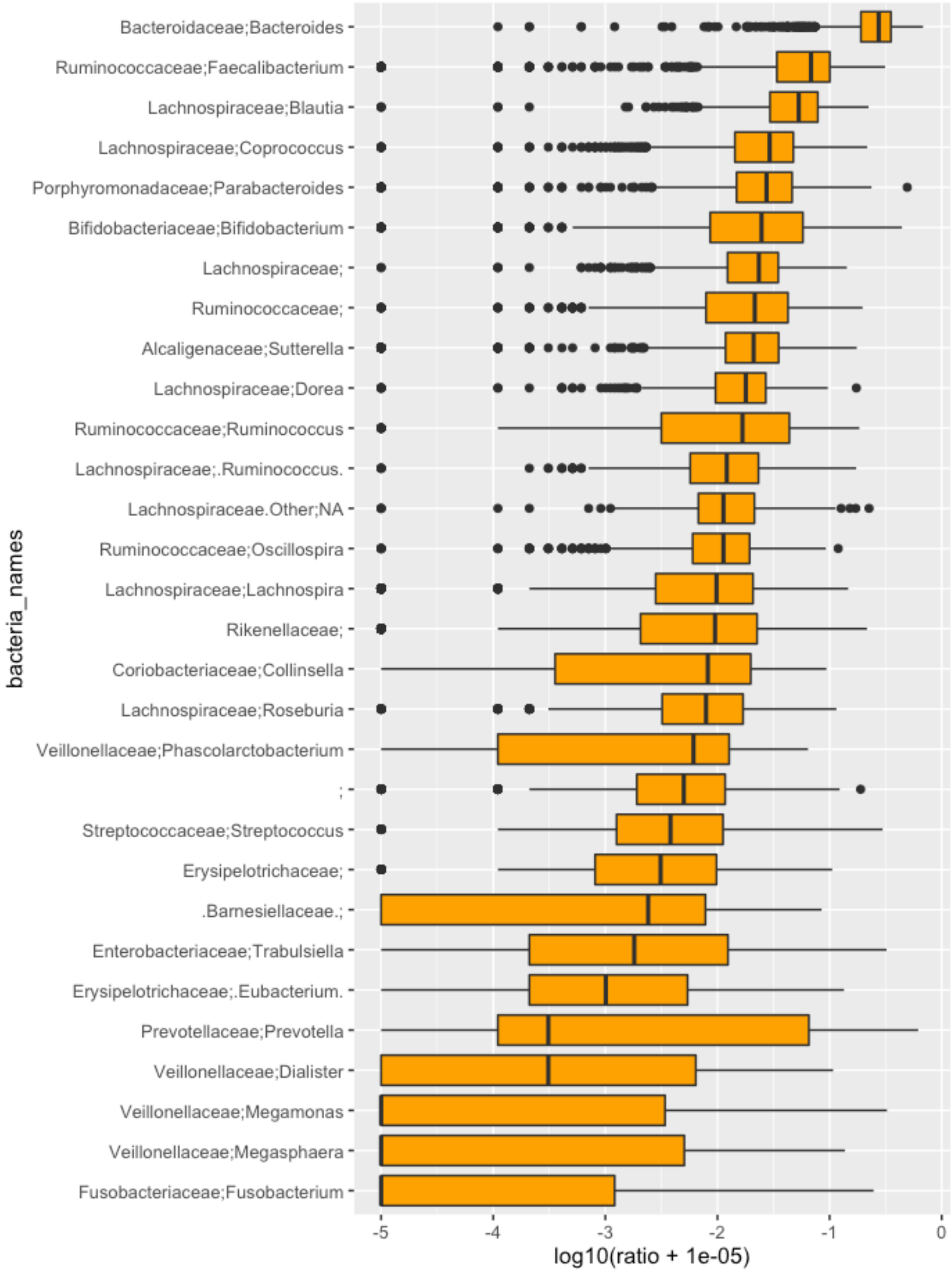


- Supplementary Figure 2
