## Supplementary Tables for "Analysis of healthy reference ranges for clinically relevant gut microbial indicators: Mykinso cohort study in Japan"

### Table S1.

| Question category | Question type | Description |
|---|---|---|
| sample_name | | The barcode for the sample |
| collection_date | | Date of sample collection |
| age_years | | The self-reported participant age in years |
| sex | | Participant biological sex, not sexual identity |
| height_cm | | Participant height in cm |
| weight_kg | | Participant weight in kilograms |
| skin_condition | | Does the participant have a skin condition? |
| food allergies | | Does the participant have food allergies? |
| seasonal_allergies | | Does the participant have seasonal allergies? |
| smoking_frequency | frequency | How often does the participant smoke? |
| alcohol_consumption | frequency | Does the participant consume alcohol? |
| alcohol_frequency | frequency | How often does the participant drink alcohol? |
| exercise_frequency | frequency | How often does the participant exercise? |
| sleep_duration | | How long does the participant sleep on an average night? |
| bowel_movement_frequency | | Number of daily bowel movements |
| bowel_movement_quality | | Does the participant tend toward constipation or diarrhea? |
| rice or bread frequency | frequency | How often does the participant eat rice or bread? |
| bran frequency | frequency | How often does the participant eat bran? |
| vegetable_frequency | frequency | How often does the participant eat vegetables? |
| fruit_frequency | frequency | How often does the participant eat fruit? |
| meat_frequency | frequency | How often does the participant eat meat? |
| fish frequency | frequency | How often does the participant eat fish? |
| eggs_frequency | frequency | How often does the participant eat eggs? |
| milk_frequency | frequency | How often does the participant drink milk? |
| cheese_frequency | frequency | How often does the participant eat cheese? |
| fermented_plant_frequency | frequency | How often does the participant eat fermented plants? |
| natto frequency | frequency | How often does the participant eat natto? |
| pickles frequency | frequency | How often does the participant eat pickles? |
| seaweed fequency | frequency | How often does the participant eat seaweed? |
| mushroom frequency | frequency | How often does the participant eat mushroom? |
| cardiovascular_disease | medical condition | Had the participant been diagnosed with cardiovascular disease? |
| gastrointestinal disease | medical condition | Had the participant been diagnosed with gastrointestinal disease? |
| liver_disease | medical condition | Had the participant been diagnosed with liver disease? |
| kidney_disease | medical condition | Had the participant been diagnosed with kidney disease? |
| osteoarticular disease | medical condition | Had the participant been diagnosed with osteoarticular disease? |
| atopic disease | medical condition | Had the participant been diagnosed with atopic disease? |
| asthma | medical condition | Had the participant been diagnosed with asthma? |
| back pain | medical condition | Had the participant been diagnosed with back pain? |
| diabetes | medical condition | Had the participant been diagnosed with diabetes? |
| hypertension | medical condition | Had the participant been diagnosed with hypertension? |
| dyslipidemia | medical condition | Had the participant been diagnosed with dyslipidemia? |
| mental_illness | medical condition | Had the participant been diagnosed with a mental illness? |
| colorectal cancer | medical condition | Had the participant been diagnosed with colorectal cancer? |
| colorectal polyp | medical condition | Had the participant been diagnosed with colorectal polyp? |
| medication history | medical condition | The self-reported participant medication history. |
| dietary supplement frequency | frequency | How often does the participant eat or drink dietary supplement? |

Table S2.

| **Years** | **Female** | **Male** | **Number of samples** |
|---|---|---|---|
| 0-9 | 34 | 34 | 68 |
| 10-19 | 46 | 32 | 78 |
| 20-29 | 514 | 222 | 736 |
| 30-39 | 932 | 603 | 1535 |
| 40-49 | 924 | 640 | 1564 |
| 50-59 | 625 | 428 | 1053 |
| 60-69 | 271 | 191 | 462 |
| 70-79 | 135 | 119 | 254 |
| 80 over | 57 | 36 | 93 |
| Sum | 3538 | 2305 | 5843 |

Table S3.

| Years | Female | Male | Number of samples |
|---|---|---|---|
| 0-9 | 31 | 31 | 62 |
| 10-19 | 18 | 21 | 39 |
| 20-29 | 110 | 51 | 161 |
| 30-39 | 177 | 87 | 264 |
| 40-49 | 184 | 89 | 273 |
| 50-59 | 171 | 72 | 243 |
| 60-69 | 95 | 55 | 150 |
| 70-79 | 75 | 59 | 134 |
| 80 over | 18 | 20 | 38 |
| Sum | 881 | 483 | 1364 |

Table S4.

| Years | Female | Male | Number of samples |
|---|---|---|---|
| 19 under | 11 | 4 | 15 |
| 20-29 | 82 | 36 | 118 |
| 30-39 | 196 | 108 | 304 |
| 40-49 | 255 | 193 | 448 |
| 50-59 | 205 | 197 | 402 |
| 60-69 | 103 | 84 | 187 |
| 70 over | 83 | 57 | 140 |
| Sum | 935 | 679 | 1614 |
